# A B-cell–cardiac autoantibody–complement axis contributes to myocarditis induced by checkpoint blockade and cardiac irradiation

**DOI:** 10.64898/2026.11.22.768334

**Authors:** Dinh Nam Tran, Shiguang Yu, Loan Thi Kim Nguyen, Alok Dwivedi, Bo Lu

## Abstract

Thoracic radiotherapy and immune checkpoint inhibitors (ICIs) improve outcomes in locally advanced lung cancer but can cause cardiac toxicity. The mechanisms by which radiation-associated cardiac injury develops into autoimmune myocarditis during ICI therapy remain incompletely defined. Using autoimmune-prone A/J mice, we investigated whether cardiac irradiation (CIR) promotes a B-cell–cardiac autoantibody–complement axis during dual PD-1 and CTLA-4 blockade. Dual ICIs plus CIR increased the incidence of myocarditis to 76.2% and cardiac fibrosis to 69.0%. CIR did not further increase global splenic B-cell activation beyond dual ICIs alone but increased cardiac troponin I-reactive B cells and IgG, circulating cardiac autoantibodies (cAAbs), myocardial IgG accumulation, and C3/C3a activation. In mice subsequently found to have grade 3–4 myocarditis on day 21, cAAbs increased by day 7 and continued to rise, preceding histologic identification of severe disease. Cobra venom factor-mediated C3 depletion reduced myocarditis incidence from 90% to 40% and fibrosis from 85% to 35%, while also decreasing B-cell activation and circulating cAAbs. In a separate LLC-OVA tumor model, C3 depletion did not produce an evident loss of the antitumor activity of combined ICIs and radiation under the conditions tested. These findings support a model in which CIR focuses ICI-associated humoral immunity toward cardiac antigens and complement amplifies cardiac injury. The early cAAb rise and protection achieved by C3 depletion provide a rationale for prospective evaluation of cAAbs as a risk marker and complement signaling as a therapeutic target.

## INTRODUCTION

Lung cancer remains a leading cause of cancer mortality worldwide (1). For patients with unresectable, locally advanced disease, immune checkpoint inhibitors (ICIs) are commonly integrated with thoracic radiation therapy (TRT) (2). Both modalities can cause cardiac injury, including myocarditis and other serious cardiovascular adverse events (3–5), and the risk may be greater in patients with pre-existing cardiovascular disease (6). Although ICI-associated myocarditis is uncommon, it can progress rapidly and is associated with substantial mortality (7). The expanding use of combined immunotherapy and TRT (8) therefore creates an urgent need to define mechanisms that distinguish direct treatment-related injury from immune-mediated cardiac damage that may persist or amplify after the initial insult. Our previous studies showed that cardiac irradiation (CIR) combined with PD-1 blockade causes cardiac injury involving both T cells (9) and B cells (10).

Most mechanistic models of ICI-associated myocarditis have focused on autoreactive T cells (11–13). However, antibodies against cardiac troponin I (cTnI) can induce cardiomyopathy in PD-1-deficient mice (14, 15), and our prior work implicated B cells and autoreactive IgG in cardiac injury after CIR plus anti–PD-1 treatment (10). Unresolved questions include whether CIR selectively amplifies cardiac antigen–directed humoral immunity during dual checkpoint blockade; whether circulating cardiac autoantibodies (cAAbs) increase before severe myocarditis is identified histologically; and whether complement is a therapeutically actionable effector of this response. Autoantibodies can accumulate in tissues, form immune complexes, and activate complement, thereby amplifying inflammatory injury (16, 17), but the contribution of an autoantibody–complement axis to myocarditis after combined CIR and dual checkpoint blockade remains undefined.

We hypothesized that CIR releases cardiac antigens that, in the setting of PD-1 and CTLA-4 blockade, promote cardiac-reactive B-cell responses and cAAbs, followed by myocardial IgG accumulation and complement activation. Using autoimmune-prone A/J mice (9, 18), we examined the temporal relationship between circulating cAAbs and subsequent histologic myocarditis, defined the myocardial IgG and complement response, and tested whether systemic C3 depletion attenuates cardiac injury. In a complementary tumor-bearing model, we assessed whether C3 depletion preserved the antitumor activity of combined treatment. Together, these experiments were designed to determine whether a B-cell–cardiac autoantibody–complement axis links CIR to ICI-associated myocarditis and represents a separable therapeutic vulnerability.

## RESULTS

### Dual ICIs and CIR increase the incidence and severity of myocarditis and cardiac fibrosis

We previously showed that cardiac irradiation (CIR) plus anti-PD-1 exacerbates cardiac injury in C57BL/6 mice (9, 23). To model clinically severe autoimmunity, we treated A/J mice with control IgG (n = 10), IgG plus CIR (n = 20), dual immune checkpoint inhibitors (ICIs) alone (n = 20), or dual ICIs plus CIR (n = 42). H&E staining showed a spectrum from no inflammation (grade 0) to mild-to-moderate (grades 1-2) and severe myocarditis (grades 3-4) (Figure 1A). Myocarditis incidence was 76.2% (95% CI, 60.5%-87.9%) with dual ICIs plus CIR, compared with 30.0% (95% CI, 11.8%-54.2%) with dual ICIs alone and 5.0% (95% CI, 0%-25.0%) with CIR alone. Severe myocarditis occurred in 28.6%, 10.0%, 5.0%, and 0% of mice in the dual ICIs plus CIR, dual ICIs, CIR, and control groups, respectively (Figure 1B). Thus, CIR substantially increased myocarditis penetrance in the setting of dual checkpoint blockade.

**Figure 1.**
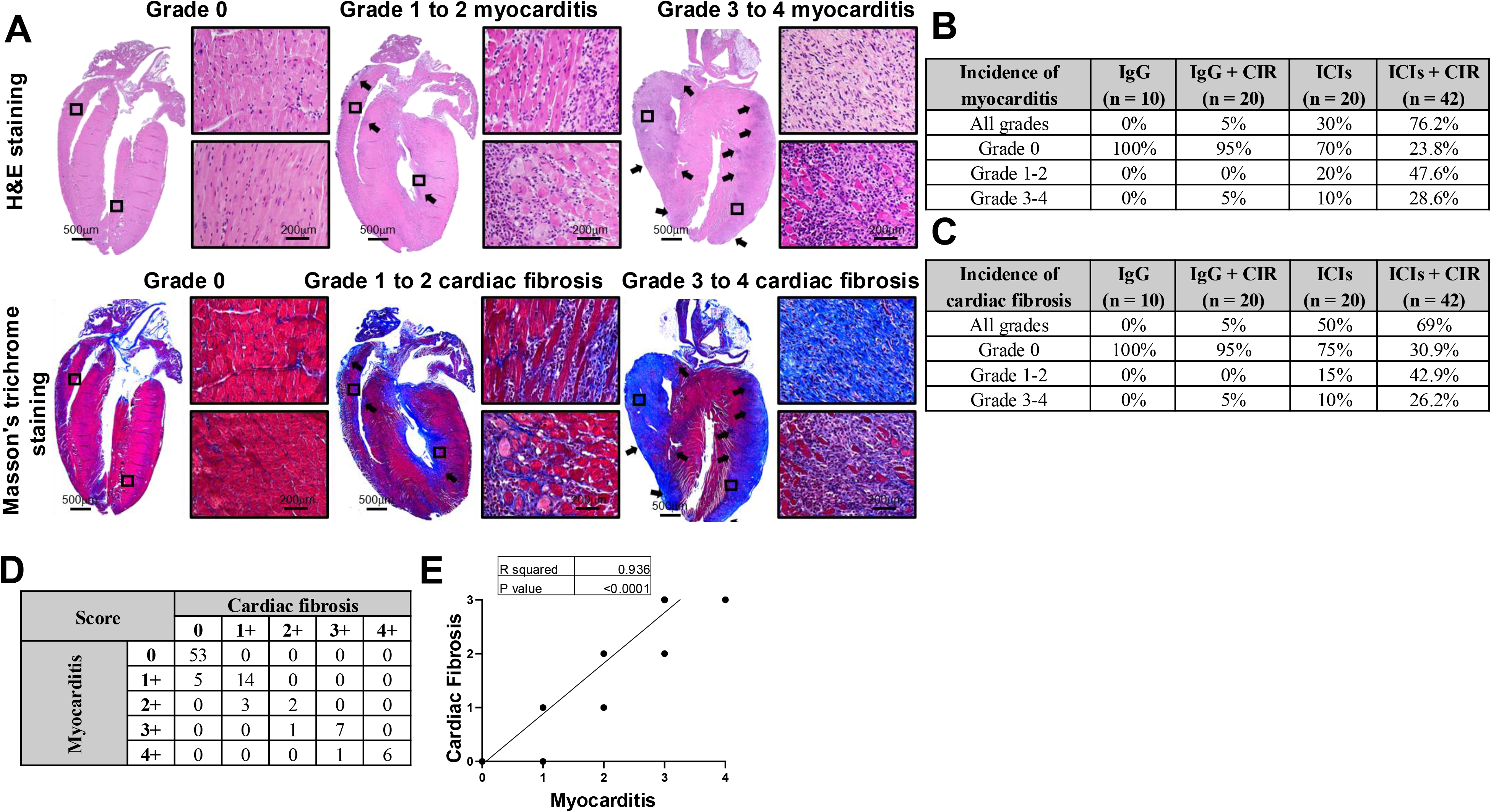
Dual immune checkpoint inhibition plus cardiac irradiation increases the incidence and severity of myocarditis and cardiac fibrosis. A, Representative hematoxylin and eosin (H&E)-stained and Masson’s trichrome-stained whole-heart sections illustrating grade 0, grades 1-2, and grades 3-4 myocarditis and cardiac fibrosis, respectively. Black arrows indicate affected regions, and boxes identify the areas shown at higher magnification. Collagen is stained blue by Masson’s trichrome. Scale bars, 500 µm (whole-heart sections) and 200 µm (higher-magnification images). B and C, Incidence and severity distributions of myocarditis and cardiac fibrosis, respectively, in mice treated with control IgG (n = 10), control IgG plus cardiac irradiation (CIR; n = 20), dual immune checkpoint inhibitors (ICIs; n = 20), or dual ICIs plus CIR (n = 42). D, Contingency table showing the relationship between myocarditis and cardiac-fibrosis grades across all animals. E, Association between myocarditis and cardiac-fibrosis grades; the reported R² and P value are shown in the panel.

Masson’s trichrome staining showed minimal collagen in normal myocardium and progressively greater fibrosis across disease grades (Figure 1A). Cardiac fibrosis was present in 69.0% (95% CI, 52.9%-82.4%) of mice treated with dual ICIs plus CIR, compared with 25.0% (95% CI, 8.7%-49.1%) treated with dual ICIs alone and 5.0% (95% CI, 0%-25.0%) treated with CIR alone. Severe fibrosis occurred in 26.2% of mice receiving dual ICIs plus CIR and 10.0% receiving dual ICIs (Figure 1C).

The contingency distribution showed that higher myocarditis grades generally occurred with higher cardiac-fibrosis grades (Figure 1D). Myocarditis and fibrosis scores were strongly associated across the observed grade combinations (R² = 0.936, P < 0.0001; Figure 1E). These findings support close coupling between myocardial inflammation and fibrotic remodeling but do not, by themselves, establish causality.

### CIR focuses ICI-driven B-cell activation toward cardiac antigens

Because our prior study implicated B cells in cardiac injury caused by CIR plus PD-1 blockade (10), we asked whether CIR increases global B-cell activation or instead amplifies responses directed toward cardiac antigens. Dual ICIs, with or without CIR, increased splenic CD86+ activated B cells and CD138+ plasmablasts or plasma cells relative to control IgG or CIR alone (P < 0.001; Figure 2A). The dual ICI and dual ICI plus CIR groups did not differ significantly, indicating that CIR did not further increase global splenic B-cell activation.

**Figure 2.**
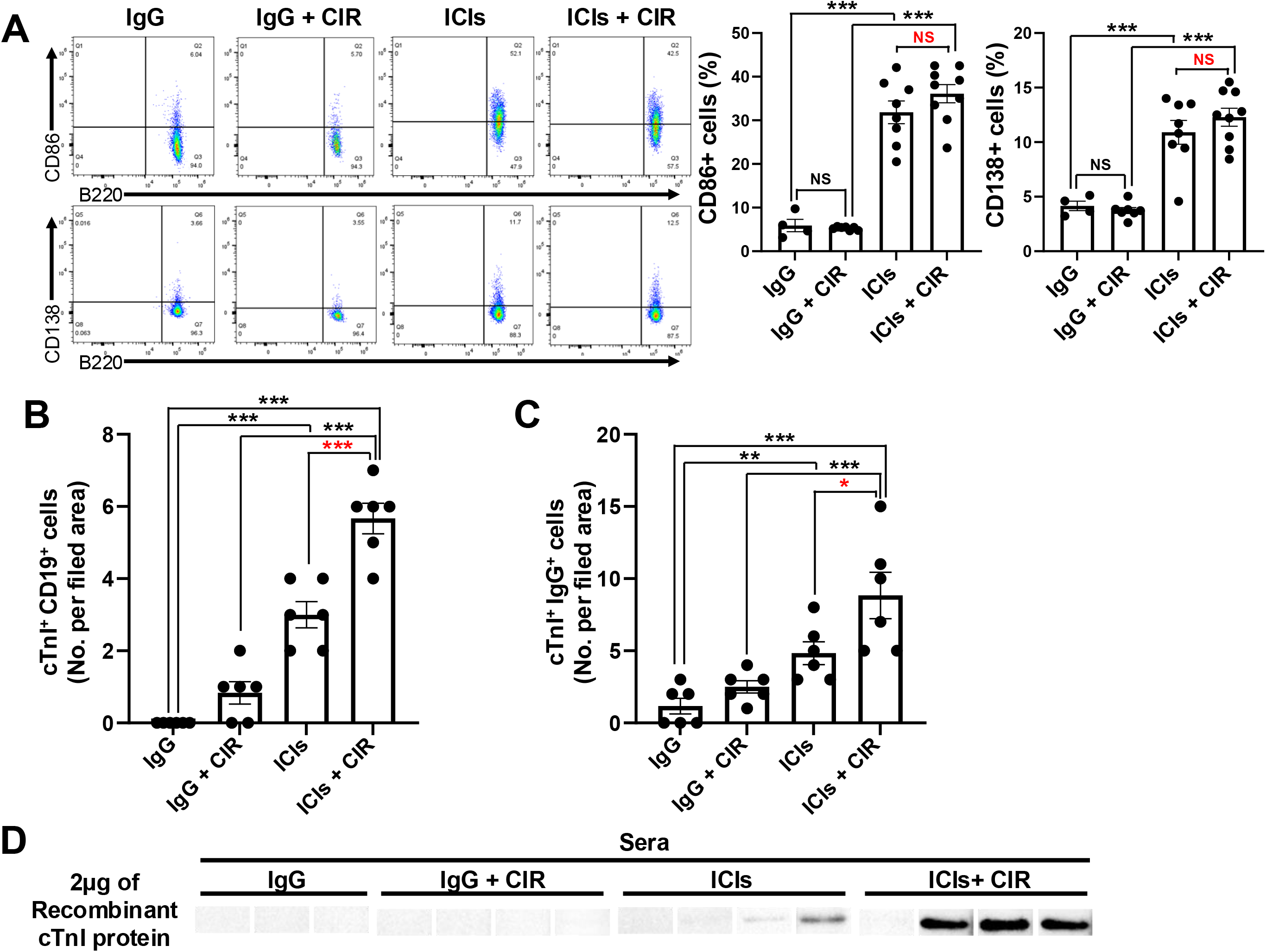
Dual immune checkpoint inhibition plus cardiac irradiation increases activated B cells, plasmablasts or plasma cells, cTnI-reactive B cells, and anti-cTnI IgG. A, Representative flow-cytometry plots and quantification of splenic CD86+ activated B cells and CD138+ plasmablasts or plasma cells in mice treated with control IgG (n = 4), control IgG plus CIR (n = 7), dual ICIs (n = 8), or dual ICIs plus CIR (n = 9). B, Quantification of cardiac troponin I (cTnI)-reactive CD19+ cells per field area in spleen (n = 6 per group). C, Quantification of cTnI-reactive IgG+ cells per field area in spleen (n = 6 per group). Representative images corresponding to panels B and C are shown in Supplementary Figure 1. D, Serum anti-cTnI IgG detected using recombinant mouse cTnI protein in mice treated with control IgG (n = 3), control IgG plus CIR (n = 4), dual ICIs (n = 4), or dual ICIs plus CIR (n = 4). Data are mean ± SEM. One-way ANOVA with Tukey’s multiple-comparison test was used. NS, not significant; *P < 0.05; **P < 0.01; ***P < 0.001.

We next assessed antigen-directed responses using cardiac troponin I (cTnI), a cardiac-restricted protein released during myocardial injury and capable of inducing experimental autoimmune myocarditis (26, 27). cTnI-reactive CD19+ cells were rare after control IgG or CIR alone, increased with dual ICIs, and were more than twofold more abundant with dual ICIs plus CIR than with dual ICIs alone (P < 0.001; Figure 2B and Supplementary Figure 1A). Splenic cTnI-reactive IgG+ cells followed the same pattern and were approximately doubled by the addition of CIR to dual ICIs (P = 0.0363; Figure 2C and Supplementary Figure 1B). Serum reactivity against recombinant cTnI was also strongest in the dual ICIs plus CIR group (Figure 2D). Together, these results distinguish broad ICI-associated B-cell activation from the cardiac antigen-directed response amplified by CIR.

### Circulating cardiac autoantibodies rise before severe myocarditis

We developed an indirect ELISA using normal mouse heart lysate to quantify serum IgG reactivity against cardiac antigens. At day 21, mice with grade 3-4 myocarditis had higher circulating cardiac autoantibody (cAAb) levels than mice with grade 0 or grades 1-2 disease (P < 0.001 for both comparisons; Figure 3A).

**Figure 3.**
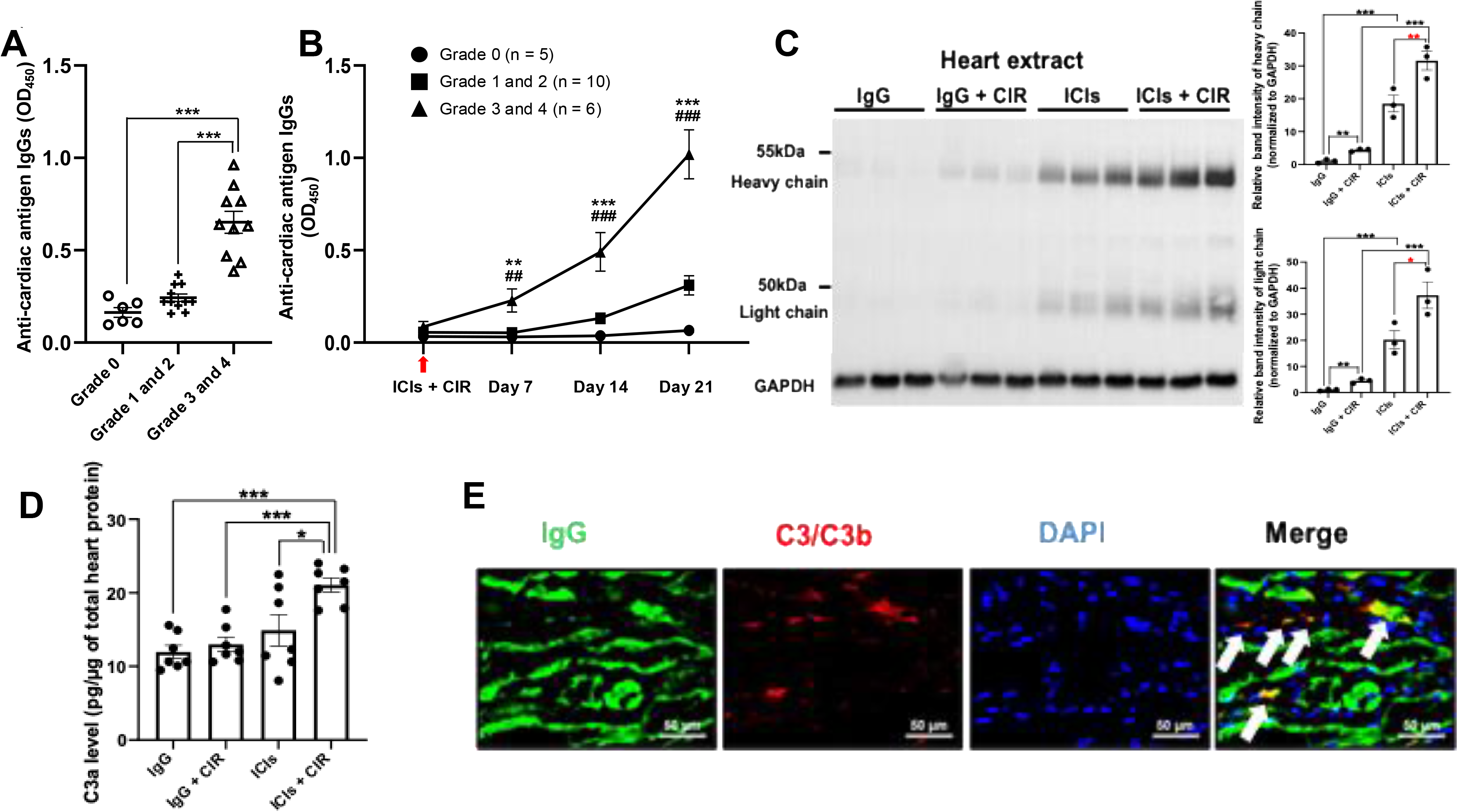
Circulating cardiac autoantibodies rise before severe myocarditis and are accompanied by myocardial IgG accumulation and complement activation. A, Circulating cardiac autoantibodies, measured by indirect ELISA and expressed as optical density at 450 nm (OD450), stratified by myocarditis grade (grade 0, n = 6; grades 1-2, n = 11; grades 3-4, n = 10). B, Serial circulating cardiac autoantibodies at baseline and on days 7, 14, and 21 after CIR in mice stratified by myocarditis severity at day 21 (grade 0, n = 5; grades 1-2, n = 10; grades 3-4, n = 6). **P < 0.01 and ***P < 0.001 versus grade 0; ##P < 0.01 and ###P < 0.001 versus grades 1-2. C, Representative immunoblot of IgG heavy and light chains in heart extracts and densitometric quantification normalized to GAPDH (n = 3 per group). D, C3a concentration in heart extracts, normalized to total heart protein. E, Representative myocardial immunofluorescence showing IgG (green), C3/C3b (red), nuclei (DAPI; blue), and merged signals. White arrows indicate IgG and C3/C3b colocalization. Scale bars, 50 µm. Data are mean ± SEM. *P < 0.05; **P < 0.01; ***P < 0.001, as indicated.

We then analyzed serial samples collected at baseline and on days 7, 14, and 21 from mice stratified by histologic severity at day 21. cAAb levels were low before treatment in all groups. In mice that subsequently developed grade 3-4 myocarditis, cAAbs increased by day 7 and continued to rise through day 21. In contrast, levels remained low in mice with grade 0 myocarditis and increased more modestly in mice with grades 1-2 myocarditis (Figure 3B). Thus, an early rise in circulating cAAbs preceded the histologic identification of severe myocarditis in this model.

### Cardiac autoantibodies are accompanied by myocardial IgG accumulation and C3 activation

To examine the effector pathway associated with the humoral response, we measured IgG in heart tissue. IgG heavy- and light-chain signals were undetectable or minimal after control IgG or CIR alone, increased with dual ICIs, and were strongest after dual ICIs plus CIR (Figure 3C).

C3a concentrations in heart extracts also increased across these treatment groups and were highest after dual ICIs plus CIR (Figure 3D). Immunofluorescence demonstrated myocardial IgG accumulation with C3/C3b deposition and areas of colocalization after dual ICIs plus CIR (Figure 3E). These findings place myocardial immunoglobulin accumulation and complement activation downstream of the treatment-associated cardiac-directed humoral response; however, colocalization alone does not establish that the deposited IgG initiated complement activation.

### C3 depletion attenuates myocarditis, fibrosis, and the associated humoral response

We next tested whether C3 activation contributes functionally to disease by administering cobra venom factor (CVF) during dual ICI plus CIR treatment. CVF reduced circulating C3 (P < 0.05; Supplementary Figure 2A), confirming target engagement, and markedly reduced myocardial C3a staining (Supplementary Figure 2B). Histologically, CVF decreased myocarditis incidence from 90% with phosphate-buffered saline (PBS) to 40%; severe myocarditis decreased from 30% to 15% (Figure 4A and 4B). Cardiac-fibrosis incidence decreased from 85% to 35%, while severe fibrosis decreased from 20% to 5% (Figure 4A and 4C).

**Figure 4.**
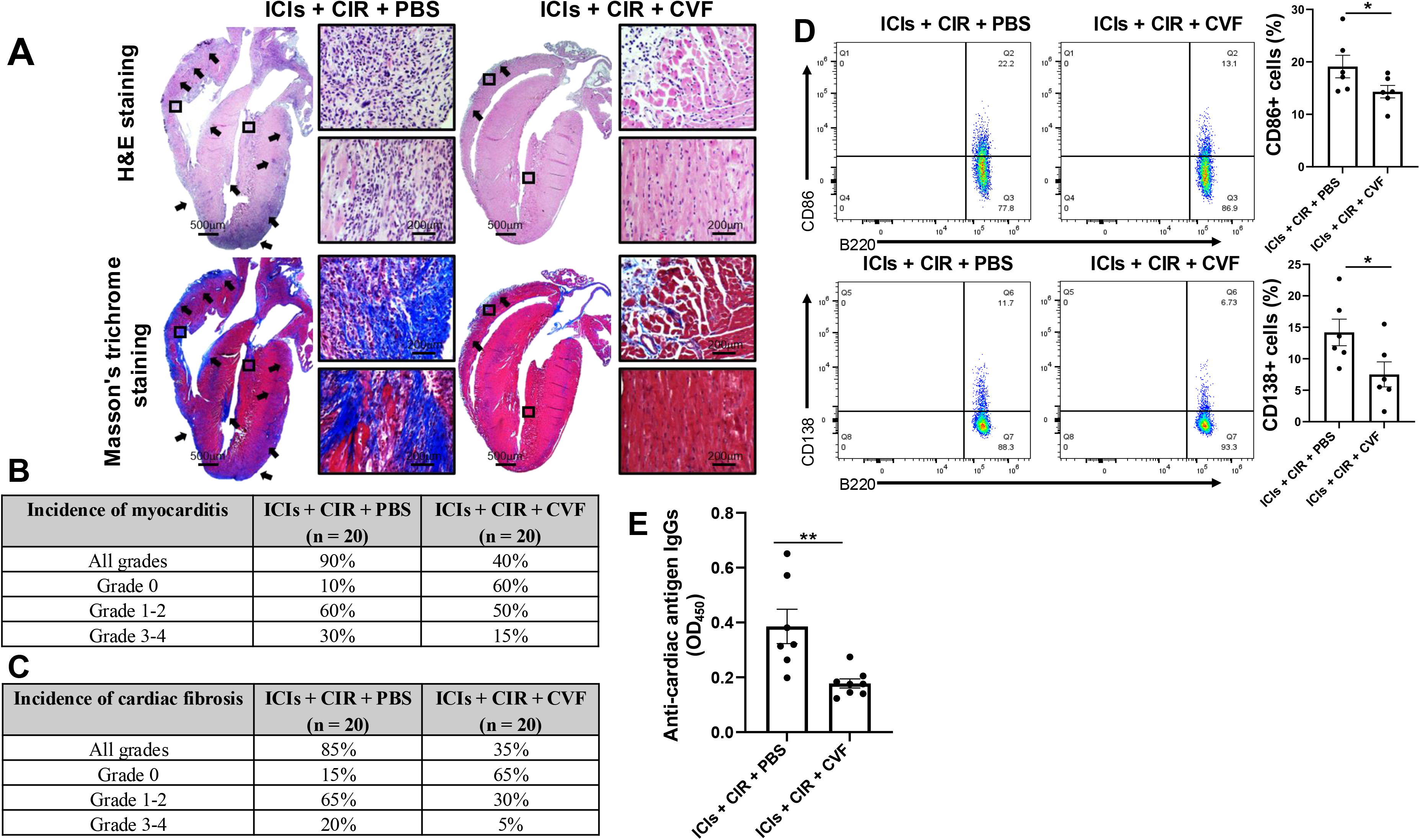
C3 depletion by cobra venom factor attenuates dual ICI plus cardiac irradiation-induced myocarditis, fibrosis, B-cell activation, and cardiac autoantibody production. A, Representative H&E-stained and Masson’s trichrome-stained whole-heart sections from mice treated with dual ICIs plus CIR and phosphate-buffered saline (PBS) or cobra venom factor (CVF). Black arrows indicate affected regions, and boxes identify the areas shown at higher magnification. Scale bars, 500 µm (whole-heart sections) and 200 µm (higher-magnification images). B and C, Incidence and severity distributions of myocarditis and cardiac fibrosis, respectively, in the PBS and CVF groups (n = 20 per group). D, Representative flow-cytometry plots and quantification of splenic CD86+ activated B cells and CD138+ plasmablasts or plasma cells (n = 6 per group). E, Serum cardiac autoantibodies measured by indirect ELISA and expressed as OD450 (PBS, n = 7; CVF, n = 8). Data are mean ± SEM. *P < 0.05; **P < 0.01, as indicated.

CVF also reduced splenic CD86+ activated B cells and CD138+ plasmablasts or plasma cells (P < 0.05 for both; Figure 4D) and lowered circulating cAAbs (P < 0.01; Figure 4E). These findings indicate that C3 depletion suppresses complement activation in the myocardium while attenuating the associated B-cell and autoantibody response.

### C3 depletion preserves the antitumor efficacy of ICIs plus radiation therapy

Because complement depletion reduced cardiac toxicity, we asked whether CVF compromised antitumor efficacy. Mice bearing subcutaneous LLC-OVA tumors were treated with vehicle, CVF, dual ICIs, radiation therapy (RT), dual ICIs plus RT, or dual ICIs plus RT and CVF. CVF did not diminish tumor-growth control achieved by dual ICIs plus 20-Gy RT and did not reduce efficacy in the 5-Gy RT regimen (Supplementary Figure 3). Thus, under the conditions tested, C3 depletion attenuated cardiac injury without an evident loss of antitumor activity.

Together, these data support a model in which CIR increases cardiac-antigen availability during dual PD-1 and CTLA-4 blockade, amplifying cardiac-reactive B-cell and autoantibody responses. Myocardial IgG accumulation and C3 activation are associated with myocarditis and fibrosis, whereas C3 depletion interrupts this axis without compromising tumor control (Figure 5).

**Figure 5.**
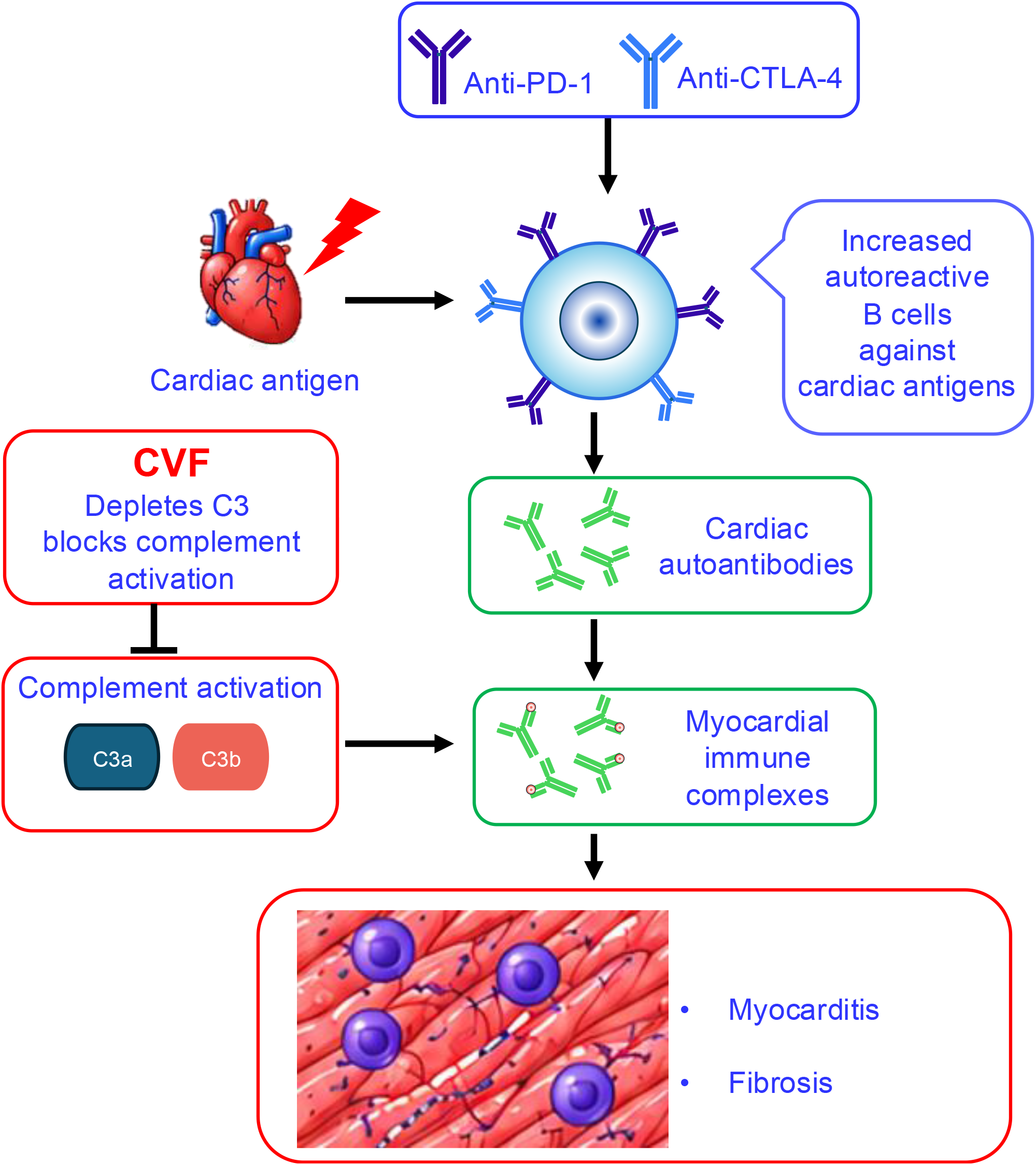
Proposed model of complement-dependent myocarditis after dual immune checkpoint inhibition and cardiac irradiation. CIR increases cardiac-antigen availability during dual PD-1 and CTLA-4 blockade, promoting cardiac-reactive B cells and cardiac autoantibodies. Myocardial immune-complex formation and complement activation, including generation or deposition of C3a and C3b, contribute to myocarditis and cardiac fibrosis. C3 depletion with CVF suppresses complement activation and attenuates these effects. The diagram summarizes the proposed relationships supported by the present experiments.

## DISCUSSION

This study identifies a treatment-associated humoral pathway linking dual immune checkpoint inhibition and cardiac irradiation (CIR) to myocarditis and cardiac fibrosis. CIR did not further increase global splenic B-cell activation beyond dual immune checkpoint inhibitors (ICIs) alone; instead, it amplified cardiac troponin I-reactive B-cell and antibody responses. Circulating cardiac autoantibodies (cAAbs) rose early in mice that were subsequently found to have severe myocarditis, while myocardial IgG accumulation was accompanied by increased C3a and C3/C3b deposition. Depletion of C3 with cobra venom factor (CVF) reduced myocarditis, fibrosis, B-cell activation, and circulating cAAbs without an evident loss of antitumor activity in a separate lung tumor model. Together, these findings support a B-cell-cardiac autoantibody-complement axis that contributes to tissue injury after combined dual ICIs and CIR.

CIR alone produced little myocarditis, fibrosis, or myocardial IgG accumulation, indicating that radiation-associated cardiac injury or antigen release was insufficient to produce the full phenotype when immune checkpoints remained intact. Dual checkpoint blockade can release autoreactive T cells from peripheral tolerance (39, 40), and CTLA-4 blockade can also shift the B-cell repertoire toward autoimmunity (41). Our findings add an antigen-directed component: although global splenic B-cell activation was similar with dual ICIs and dual ICIs plus CIR, cTnI-reactive CD19+ cells, cTnI-reactive IgG+ cells, serum reactivity against recombinant cTnI, and broader cAAbs were increased when CIR was added. We therefore propose that dual ICIs create a permissive immune state, whereas irradiated cardiac tissue supplies or exposes antigens that focus the humoral response. Direct lineage tracing, B-cell receptor repertoire analysis, and identification of the dominant antibody specificities will be required to establish this sequence.

Myocarditis and cardiac-fibrosis grades were strongly associated at the terminal assessment. This relationship is biologically consistent with inflammatory cardiomyocyte injury and reparative fibrotic remodeling (42–47), although our results are insufficient to support a causal relationship. The reduction of both myocarditis and fibrosis after C3 depletion is consistent with complement acting upstream of, or contributing to, tissue remodeling; however, the current study does not define the cellular intermediates or temporal transition from inflammation to fibrosis.

The temporal cAAb analysis provides an important distinction between an early circulating signal and an endpoint correlate. In mice ultimately classified with grade 3-4 myocarditis, cAAbs increased by day 7 and continued to rise through day 21, whereas levels remained low in mice with grade 0 disease and increased more modestly in mice with grades 1-2 disease. Because cardiac histology was obtained only on day 21, these data show that the cAAb rise preceded histologic identification of severe myocarditis, not necessarily the biological onset of myocardial inflammation. cAAbs should therefore be considered a candidate dynamic risk marker in this model rather than an established diagnostic biomarker. Prospective clinical studies should pair serial cAAb measurements with troponin, electrocardiography, cardiac imaging, treatment timing, and adjudicated myocarditis criteria.

The tissue and intervention findings strengthen the mechanistic interpretation while defining its boundaries. Myocardial IgG heavy- and light-chain abundance and C3a concentration were greatest after dual ICIs plus CIR, and immunofluorescence demonstrated areas of IgG and C3/C3b colocalization. These data establish concurrent myocardial immunoglobulin accumulation and complement activation but do not prove that the deposited IgG initiated complement activation or that immune complexes were directly formed. CVF reduced circulating C3, myocardial C3a staining, circulating cAAbs, and splenic CD86+ and CD138+ B-cell populations in parallel with histologic protection. Because systemic C3 depletion can affect both complement effector activity and B-cell responses, the present experiments do not separate these mechanisms. The reduction in B-cell and autoantibody readouts is consistent with the ability of complement-receptor signaling to amplify humoral immunity (65), but cell-specific studies will be needed to determine the responsible subtypes of immune cells. These findings complement rather than replace established T-cell and myeloid mechanisms of ICI-associated myocarditis (67–70). A plausible integrated model is that checkpoint blockade permits autoreactive lymphocyte activation, CIR increases cardiac-antigen availability, cardiac-directed antibodies accumulate in the myocardium, and complement amplifies inflammatory injury and fibrosis (Figure 5). The relative contribution of this axis will likely vary with host genetics, radiation exposure, ICI regimen, and disease stage.

Complement-directed therapy may offer an advantage over broad B-cell depletion by targeting an inflammatory amplifier while potentially preserving immune functions required for tumor control and tissue homeostasis (71, 72). In the separate subcutaneous LLC-OVA model, adding CVF did not diminish tumor-growth control achieved by dual ICIs plus 20-Gy radiation therapy and did not reduce activity in the 5-Gy regimen. This finding is encouraging but model-specific. CVF is an experimental and broadly acting complement-depleting reagent, the tumor model was subcutaneous, group sizes were modest, and follow-up was limited to 11 days. Moreover, complement can support radiation-induced antitumor immunity in some settings (73–75). The optimal component, timing, and degree of complement inhibition will therefore require testing in orthotopic and immunologically diverse tumor models.

Our studies were designed for mechanistic study. We favored a mouse with highest incidence of myocarditis at high grades by treating male autoimmune-prone A/J mice with a 20-Gy cardiac irradiation and dual ICIs, which is far beyond heterogeneous cardiac exposure encountered during contemporary thoracic radiotherapy used to treat lung cancer patients. Our tissue analyses were mostly performed around three weeks after CIR, which detects the acute phase of cardiac injury as demonstrated by tissue inflammation, which is exacerbated in the presence of dual ICIs. We acknowledge that CVF is not a clinically selective complement inhibitor and plan to test the efficacy of clinically tested complement inhibitors. Our ongoing studies evaluate potential predictive and diagnostic performance of cAAbs detected in the blood samples from lung cancer patients with autoimmune cardiotoxicities due to thoracic radiotherapy and ICI-based immunotherapy. These studies will validate the temporal cAAb signal in larger and prospectively sampled animal cohorts and in patients receiving ICIs with or without thoracic radiotherapy. We aim to identify the dominant cardiac antigens and antibody clonotypes; determine whether cAAbs are pathogenic, prognostic, or both; directly characterize myocardial immune complexes; and distinguish complement effects on B cells from complement-mediated tissue injury.

In conclusion, combined dual ICIs and CIR engage a cardiac-directed B-cell and autoantibody response associated with myocardial complement activation. The early rise of circulating cAAbs and the protection achieved by C3 depletion support prospective evaluation of cAAbs as a risk marker and complement signaling as a therapeutic target. These findings provide mechanistic and translational rationale for further investigation but do not establish cAAbs as a diagnostic test or complement inhibition as standard treatment.

## METHODS

### Mice

All mouse experiments were approved by the Institutional Animal Care and Use Committee at the University of Missouri. Eight-week-old male A/J mice were used for the myocarditis experiments, as previously described (18). Male C57BL/6 mice were used for the lung cancer experiment (19). Both strains were obtained from The Jackson Laboratory (Bar Harbor, ME, USA) and randomly assigned to experimental groups. Mice were housed in a designated animal care facility, and all procedures conformed to the Guide for the Care and Use of Laboratory Animals (8th ed., National Academies Press, 2011).

### Mouse model of myocarditis

Experiments were conducted in male mice because this model develops myocarditis at a higher incidence and severity in males than in females (20). A/J mice received 150 µg of control IgG (catalog no. BE0089, clone 2A3; Bio X Cell, Lebanon, NH, USA) or 150 µg each of anti-PD-1 (catalog no. BE0146, clone RMP1-14; Bio X Cell) and anti-CTLA-4 (catalog no. BP0032, clone UC10-4F10-11; Bio X Cell) in 100 µL phosphate-buffered saline (PBS; catalog no. 10-010-03; Thermo Fisher Scientific, Waltham, MA, USA). Treatment began 1 day before cardiac irradiation (CIR) and was repeated every 3 days on days 2, 5, 8, 11, 14, 17, and 20. Antibody doses were selected as previously described (18).

CIR was delivered on day 0 using a Precision X-RAD SmART irradiator (Precision X-Ray, North Branford, CT, USA), with planning to minimize lung and esophageal exposure. A total dose of 20 Gy was delivered at 225 kV and 20 mA using a 0.3-mm copper filter and a 10-mm C10 collimator; control mice received 0 Gy. On day 21, mice were euthanized and perfused with cold PBS. Hearts were fixed for histology and immunofluorescence or snap-frozen and stored at -80°C for protein extraction. Spleens and serum were collected for cellular and antibody analyses.

### C3 depletion with cobra venom factor

To deplete C3, mice receiving dual ICIs plus CIR were treated intraperitoneally with cobra venom factor (CVF; catalog no. 233552; Sigma-Aldrich, St. Louis, MO, USA) at 1 unit in 150 µL sterile PBS or with PBS vehicle. Treatment began 1 day before CIR and continued on days 1, 2, 3, 6, 7, 8, 10, 11, 13, 14, 16, 19, and 20 after CIR, as previously described (21). Animals were euthanized on day 21, and tissues and serum were collected as described above.

### Histological evaluation of myocarditis and cardiac fibrosis

Heart tissues were fixed in 10% buffered paraformaldehyde for 12 hours at 4°C, embedded in paraffin, and sectioned at 4 µm. Hematoxylin and eosin (H&E) staining was used to assess myocarditis, as previously described (18). An inflammatory infiltrate was defined morphologically as a discrete interstitial or perivascular aggregate of mononuclear nucleated cells separating or replacing cardiomyocytes, with or without adjacent myocyte injury. Two investigators blinded to treatment graded myocarditis according to the proportion of the examined section occupied by inflammatory infiltrates: grade 0, none; grade 1, <10%; grade 2, 10%-35%; grade 3, 35%-50%; and grade 4, >50%. H&E morphology was not used to assign a specific immune-cell lineage. Cardiac fibrosis was assessed using Masson’s trichrome staining (KTMTR Trichrome Stain Kit-Masson’s; StatLab, McKinney, TX, USA) according to the manufacturer’s instructions. Collagen-positive tissue was identified by blue staining. Fibrosis was graded using the same percentage thresholds as the myocarditis scale, based on the proportion of the examined section occupied by collagen-positive tissue. Images were analyzed using ImageJ (National Institutes of Health, Bethesda, MD, USA).

### Identification of cTnI-reactive B cells and IgG

Spleens were collected on day 21 and stored at -80°C. Frozen sections were fixed in 2% paraformaldehyde for 20 minutes, washed with PBS, and blocked with 5% goat serum and Fc blocker (catalog no. S17011E; BioLegend, San Diego, CA, USA) for 1 hour. Sections were incubated overnight at 4°C with recombinant His-tagged cardiac troponin I (cTnI; 2 µg/mL; catalog no. ab316587; Abcam, Cambridge, UK). Alexa Fluor 594-conjugated rabbit anti-His-tag antibody (catalog no. C39361-AF594; Signalway Antibody, Greenbelt, MD, USA) was used to detect cTnI. Mouse IgG was detected with Alexa Fluor 488-conjugated goat anti-mouse IgG (1:5000; catalog no. A-11001; Invitrogen/Thermo Fisher Scientific). For CD19 staining, sections were incubated overnight at 4°C with anti-CD19 (1:250; catalog no. ab245235; Abcam), washed, and incubated for 1 hour with Alexa Fluor 488-conjugated secondary antibody (catalog no. A28175; Invitrogen). Nuclei were counterstained with DAPI (catalog no. 50-874-10001; MilliporeSigma, St. Louis, MO, USA). Sections were imaged at 40× using a Leica confocal microscope (Buffalo Grove, IL, USA). cTnI-reactive CD19+ and cTnI-reactive IgG+ signals were counted per imaged field area in matched splenic regions; group analyses included six mice per treatment condition.

### Serum anti-cTnI IgG assay

Serum anti-cTnI IgG was assessed using recombinant mouse cTnI protein. Equal quantities of recombinant cTnI (2 µg) were prepared for each serum lane, and serum samples were applied at the same dilution. Bound mouse IgG was detected with an anti-mouse IgG secondary antibody. Because the assay did not include a validated serum loading or total-IgG normalization control, the resulting bands were interpreted qualitatively. Quantitative circulating cardiac-reactive IgG was assessed separately using the cardiac-lysate ELISA described below.

### Flow cytometry

Spleens were collected on day 21 and dissociated into single-cell suspensions, which were filtered through 70-µm strainers. Red blood cells were lysed according to the manufacturer’s instructions. After Fc-receptor blockade, cells were stained with FITC anti-mouse/human CD45R/B220 (catalog no. 103206), Alexa Fluor 647 anti-mouse CD21/CD35 (catalog no. 123424), PE anti-mouse CD23 (catalog no. 101608), Brilliant Violet 605 anti-mouse CD86 (catalog no. 105037), and Brilliant Violet 421 anti-mouse CD138 (catalog no. 142523; all from BioLegend). Samples were acquired on a Cytek Aurora and analyzed using FlowJo version 10 (BD Biosciences, Franklin Lakes, NJ, USA). CD86+ and CD138+ populations were quantified among B220+ cells.

### Myocardial IgG immunoblotting

Heart proteins were extracted in lysis buffer containing 10 mM Tris-HCl (pH 7.4), 150 mM NaCl, 2.5 mM EDTA, and 0.125% Nonidet P-40 supplemented with protease and phosphatase inhibitor cocktails. Ten micrograms of protein per sample were separated on 8% SDS-PAGE gels and transferred to polyvinylidene difluoride membranes. Membranes were blocked with 5% skim milk in PBS and probed with anti-mouse IgG (1:5000; catalog no. 31431; Invitrogen). GAPDH (1:1000; catalog no. 97166S; Cell Signaling Technology, Danvers, MA, USA) served as the loading control. Immunoreactivity was detected by enhanced chemiluminescence. IgG heavy- and light-chain band intensities were quantified using ImageJ and normalized to GAPDH.

### Measurement of C3, C3a, and myocardial C3/C3b

Serum C3 was measured using a C3-specific ELISA according to the manufacturer’s instructions. C3a in heart extracts was measured by ELISA and normalized to total heart protein. Samples were analyzed using the same acquisition and background-subtraction procedures within each assay.

For myocardial immunofluorescence, frozen heart sections were fixed in 2% paraformaldehyde for 20 minutes, rinsed with PBS, and blocked with 5% goat serum (catalog no. 31872; Thermo Fisher Scientific) and Fc blocker (1:500; catalog no. S17011E; BioLegend) for 1 hour. Mouse IgG was detected with Alexa Fluor 488-conjugated goat anti-mouse IgG (1:5000; catalog no. A-11001; Invitrogen/Thermo Fisher Scientific). C3/C3b and C3a were detected using anti-C3/C3b (1:250; catalog no. ab322300; Abcam) and anti-C3a (1:100; catalog no. ABIN7438601; Antibodies Online, Limerick, PA, USA), respectively, followed by Alexa Fluor 594-conjugated goat anti-rabbit IgG Fab fragment (1:5000; catalog no. A11072; Invitrogen). Nuclei were counterstained with DAPI. Sections were imaged at 40× using a Leica confocal microscope.

### Detection of circulating cardiac autoantibodies

Circulating cardiac autoantibodies (cAAbs) were quantified by indirect ELISA using cardiac lysate from untreated A/J mice. Heart tissue was homogenized in 1 mL radioimmunoprecipitation assay buffer supplemented with protease inhibitor. Ninety-six-well plates were coated overnight at 4°C with mouse cardiac lysate (5 µg/mL) in carbonate coating buffer. After washing with PBS containing 0.05% Tween 20 (PBST), plates were blocked with 5% bovine serum albumin in PBST for 1 hour at room temperature. Serum samples were diluted 1:1000 in blocking buffer and incubated for 2 hours at room temperature. After washing, horseradish peroxidase-conjugated anti-mouse IgG (H+L; 1:4000; catalog no. 31431; Invitrogen) was added for 1 hour. Plates were developed with TMB substrate (catalog no. 34021; Thermo Fisher Scientific), and the reaction was stopped with 2 N sulfuric acid. Absorbance was measured at 450 nm. Samples were analyzed in duplicate, and blank-well absorbance was subtracted. Values are reported as background-subtracted OD450 because no cardiac-antigen-specific reference standard was available.

For longitudinal analyses, blood was collected at baseline and on days 7, 14, and 21. Mice were stratified retrospectively according to myocarditis severity determined histologically on day 21.

### Lung tumor efficacy experiment

Male C57BL/6 mice bearing subcutaneous LLC-OVA tumors were studied using the previously described model (19, 22). When tumors reached approximately 80-100 mmS, mice were assigned to vehicle (n = 8), CVF alone (n = 8), dual ICIs alone (n = 8), radiation therapy (RT) at 20 Gy (n = 8), dual ICIs plus RT at 20 Gy (n = 8), dual ICIs plus RT at 20 Gy and CVF (n = 8), dual ICIs plus RT at 5 Gy (n = 6), or dual ICIs plus RT at 5 Gy and CVF (n = 7). ICIs with or without CVF were initiated on experimental day 0, and tumor-directed RT was delivered according to the treatment timeline. Tumor dimensions were measured serially, and tumor volume was followed through day 11. This experiment was conducted separately from the A/J myocarditis experiments.

### Statistical analysis

Categorical data are summarized as counts and percentages, and continuous data as mean ± SEM. Myocarditis and fibrosis incidence and 95% CIs were estimated using the binomial distribution and compared across groups using Fisher’s exact test. Continuous outcomes were compared between two groups using two-tailed unpaired t tests and among more than two groups using one-way analysis of variance followed by Tukey’s multiple-comparison test. Longitudinal cAAb measurements were analyzed using repeated-measures analysis of variance with multiple comparisons. The relationship between myocarditis and cardiac-fibrosis grades was evaluated by linear regression, with R² reported as the measure of association. Analyses were performed using GraphPad InStat and Stata version 19. All P values were two-sided, and P < 0.05 was considered statistically significant.

## Acknowledgments

The authors thank Dr Xiaofan Wang (Department of Pharmacology and Cancer Biology, Duke University) for providing the LLC-OVA cell line; Ms Nhan Vu for technical support; and Mr Eric Zhu and Mr Bennett Flannagan for comments and proofreading.

## Author contributions

D.N.T.: conceptualization, data curation, formal analysis, validation, investigation, visualization, methodology, writing—original draft, and writing—review and editing. S.Y.: conceptualization, data curation, formal analysis, methodology, writing—original draft, and writing—review and editing. L.T.K.N.: data curation, formal analysis, and writing—review and editing. A.D.: formal analysis and writing—review and editing. B.L.: conceptualization, resources, supervision, funding acquisition, project administration, and writing—review and editing.

## Data availability statement

The raw data generated in this study are available from the corresponding author on reasonable request.

## Nonstandard Abbreviations and Acronyms

TRT: thoracic radiotherapy
ICIs: immune checkpoint inhibitors
CTLA-4: cytotoxic T-lymphocyte-associated protein 4
PD-1: programmed cell death protein 1
dual ICIs: anti-PD-1 plus anti-CTLA-4
CIR: cardiac irradiation
cTnI: cardiac troponin I
CVF: cobra venom factor
irAEs: immune-related adverse events
cAAbs: cardiac autoantibodies

## SUPPLEMENTARY FIGURE LEGENDS

**Supplementary Figure 1.**
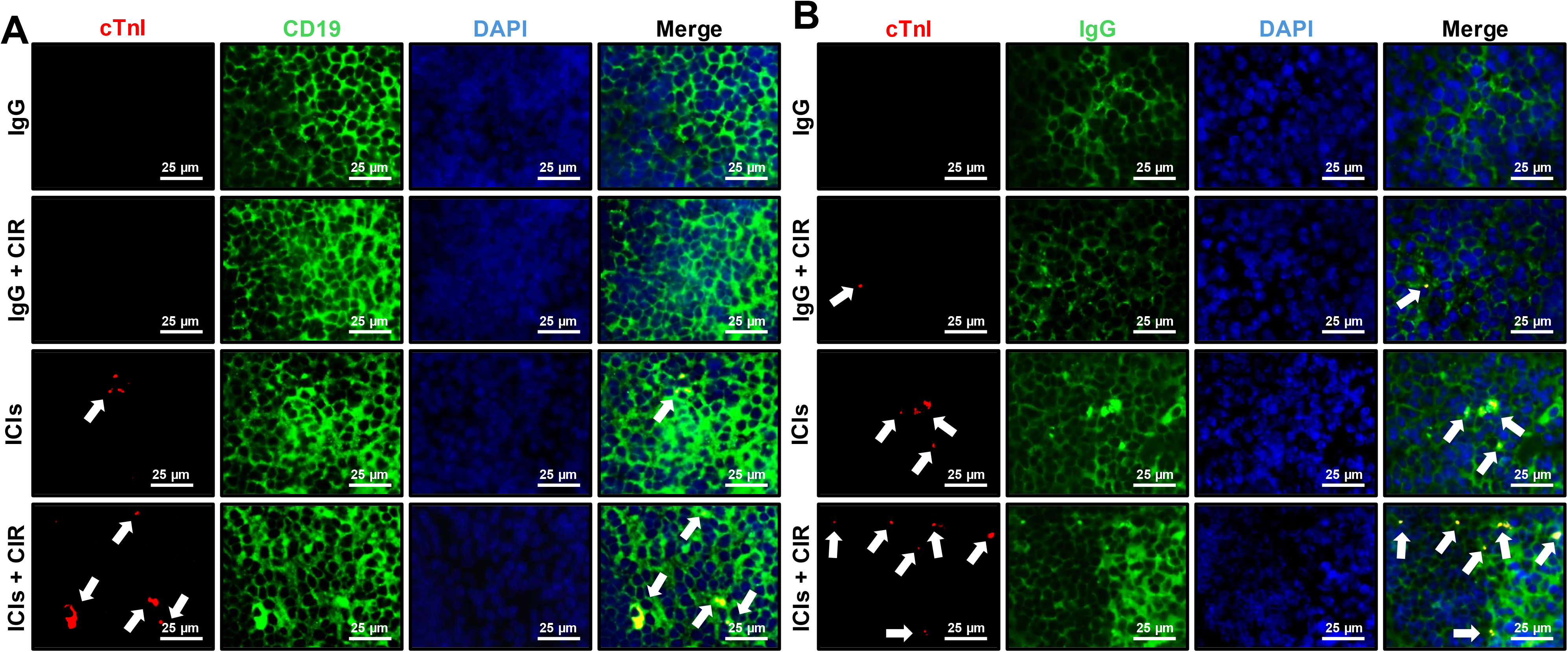
Dual immune checkpoint inhibition plus cardiac irradiation increases cTnI-reactive CD19+ cells and cTnI-reactive IgG+ cells in spleen. A, Representative immunofluorescence images of cTnI (red), CD19 (green), nuclei (DAPI; blue), and merged signals in mice treated with control IgG, control IgG plus CIR, dual ICIs, or dual ICIs plus CIR. B, Representative immunofluorescence images of cTnI (red), IgG (green), nuclei (DAPI; blue), and merged signals in the same treatment groups. White arrows indicate colocalized signals. Scale bars, 25 µm. Quantification is shown in Figure 2B and 2C.

**Supplementary Figure 2.**
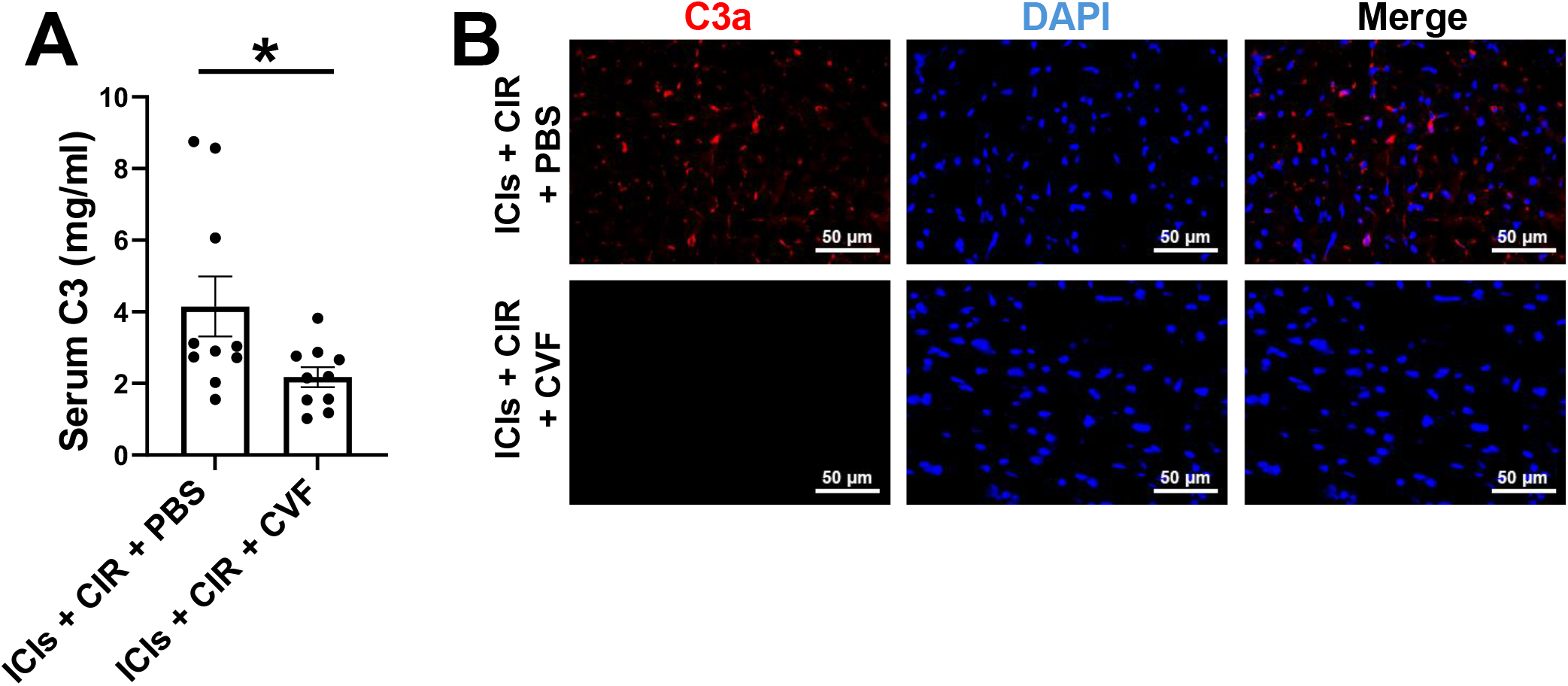
Cobra venom factor reduces circulating C3 and myocardial C3a after dual immune checkpoint inhibition plus cardiac irradiation. A, Serum C3 concentrations measured by C3-specific ELISA in mice treated with dual ICIs plus CIR and PBS or CVF. B, Representative myocardial immunofluorescence showing C3a (red), nuclei (DAPI; blue), and merged signals in the PBS and CVF groups. Scale bars, 50 µm. Data are mean ± SEM. *P < 0.05 by two-tailed unpaired t test.

**Supplementary Figure 3.**
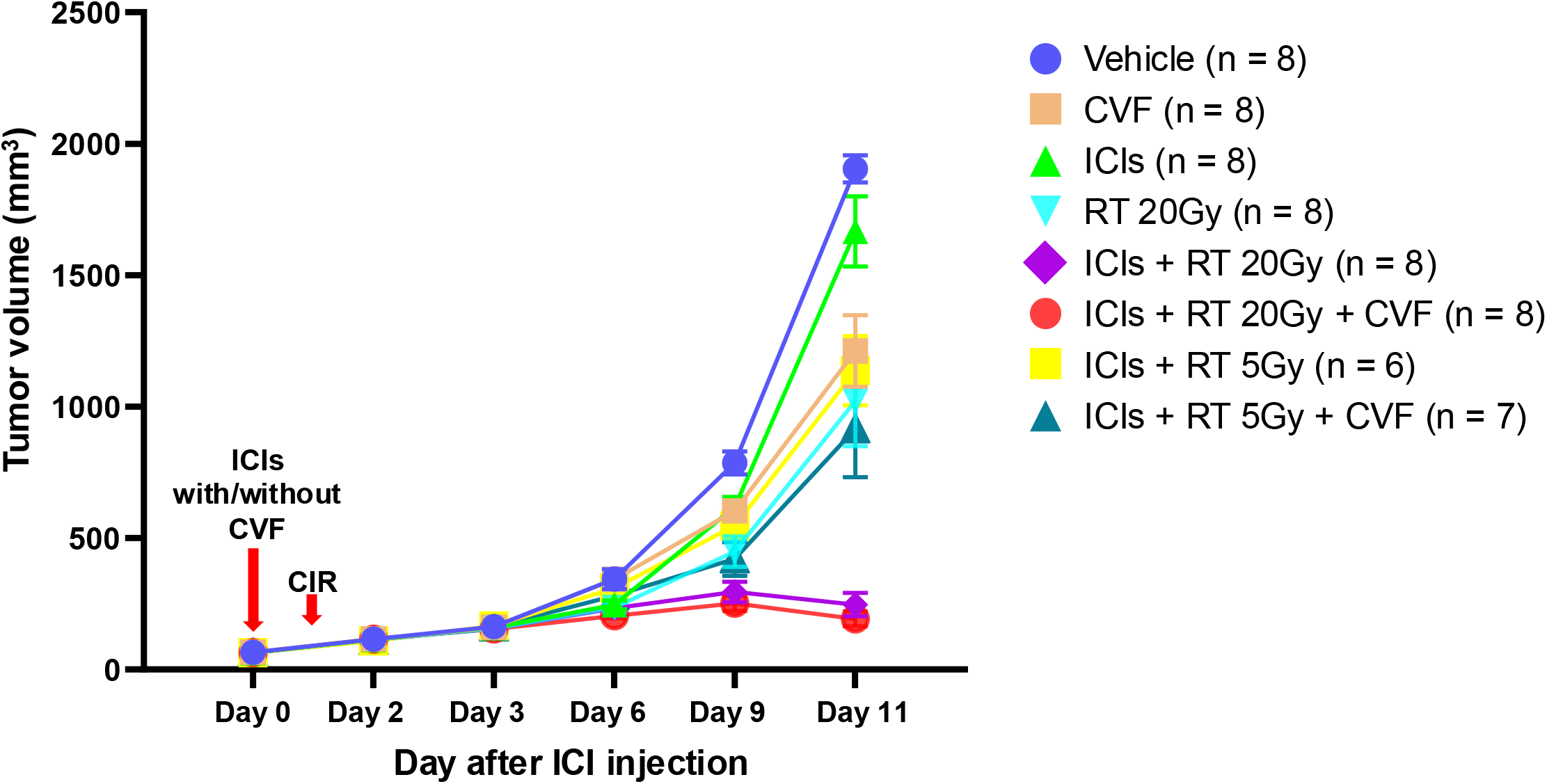
C3 depletion preserves the antitumor effects of immune checkpoint inhibition and radiation therapy. Tumor-growth curves for mice treated with vehicle (n = 8), CVF (n = 8), dual ICIs (n = 8), radiation therapy (RT) at 20 Gy (n = 8), dual ICIs plus RT at 20 Gy (n = 8), dual ICIs plus RT at 20 Gy and CVF (n = 8), dual ICIs plus RT at 5 Gy (n = 6), or dual ICIs plus RT at 5 Gy and CVF (n = 7). The treatment timeline indicates administration of ICIs with or without CVF and CIR. Data are mean ± SEM.

